# Understanding Microbeads Stacking in Deformable Nano-Sieve for Efficient Plasma Separation and Blood Cell Retrieval

**DOI:** 10.1101/2021.05.13.444053

**Authors:** Xinye Chen, Shuhuan Zhang, Yu Gan, Rui Liu, Ruo-Qian Wang, Ke Du

## Abstract

Efficient separation of blood cells and plasma is key for numerous molecular diagnosis and therapeutics applications. Despite various microfluidics-based separation strategies have been developed, a simple, reliable, and multiplexing separation device that can process a large volume of blood is still missing. Here we show a microbead packed deformable microfluidic system that can efficiently separate highly purified plasma from whole blood as well as retrieve blocked blood cells from the device. Combining microscope imaging, optical tomography scanning, and computational fluidic modeling, a highly accurate model is constructed to understand the link between the mechanical properties of the microfluidics, flow rate, and microbeads packing/leaking, which supports and rationalizes the experimental observations. This deformable nano-sieve device establishes a key technology for centrifuge-free diagnosis and treatment of bloodborne diseases and may be important for the design of next-generation deformable microfluidics for separation applications.

## Introduction

Blood is a multi-phase biological fluid that can be completely analyzed to present the fundamental functions of a human body, such as homeostasis [1], nutrients transportation [2], regulation of body temperature and pH [3], circulation of oxygen [4], and immune cells throughout the body [5]. Cell components and protein-enriched plasma are two major components in blood. The cells suspension consists of white blood cells, red blood cell (RBCs), and platelets. The analysis of physical properties of cell components in blood can be used for the diagnosis of several cell-based diseases, such as cancer [6,7], sepsis [8,9], sickle-cell anemia [10,11], and malaria [12,13]. On the other hand, human plasma that is clear and straw-colored liquid (54.3% by volume of whole blood) comprises various proteins (e.g., fibrinogen, globulin, and albumin) and several inorganic compounds [14]. Efficiently separating plasma from whole blood is the key for molecular diagnostics and therapeutics. For instance, testing the relative proportions of biological substances from the plasma, e.g., cholesterol and glucose, can be used to evaluate the health condition of internal organs of a human body [5]. The quantitative analysis of specific protein biomarkers in plasma can be used to diagnose many diseases such as genetic abnormalities [15,16], paraproteinemias [17,18], and hemoglobinopathies [19,20]. Therefore, extracting highly purified plasma from whole blood is important to the precise assessment and diagnosis of those diseases.

Conventionally, extracting human plasma from whole blood in laboratories and clinics relies on centrifugation process, requiring expensive and bulky instrumentations, skilled technician, and time-consuming protocols. The throughput is low as only a few samples can be processed at the same time. On the other hand, various microfluidic platforms have been introduced to separate plasma from whole blood. For example, well-defined weir structures with a gap size of 0.5 μm have been demonstrated to continuously separate the plasma in a cross-flow configuration [21]. The small gap can block the cell components and allows the cell-free plasma to flow through the microfluidic channel. Similarly, a transverse-flow microfilters with the same gap size was reported to efficiently separate plasma [22]. Moorthy *et al*. developed a novel filter with porous structures in a microchannel by an emulsion photo polymerization process [23]. However, the fabrication and integration of sub-micron features into microfluidic devices are challenging. The small fluidic passage also limits the filtration speed. In addition, due to the high fluidic resistance, blood needs to be extensively diluted (typically 1:20), further delaying the processing time. Recently, an alternative method based on microbead-stacked microchannel has been developed [24]. This microbead-based filter impeded the RBCs transportation and permitted the cell-free plasma to drive through the microbead array by capillary force without dilution. However, the slow capillary action significantly increased the processing time and is not suitable when dealing with a large volume of blood. Therefore, developing a miniaturized and simple microfluidic device that can be used for multiplexing and rapid plasma separation from whole blood is urgently needed for various biomedical applications [25].

However, designing a microfluidic device that can separate various biomarkers is challenging because the operation of the device requires a good understanding of the device behavior and the link between theory and practice has been under-developed. A seminal work by Christov *et al*. developed a theoretical model to establish the relationship between flow rate and microfluidic deformation, which has been found comparing well with the experiments [26]. We further extended this model and applied it to the design of deformable microfluidics and showed good fitting with the experimental observation [27]. However, those models only concentrated on the change of the flow properties and less is known on how the properties of the structure change will impact the nano-sieve performance and to what extent the model will be valid to apply.

In this study, we present a deformable microbead stacked nano-sieve system for efficient plasma separation from whole blood, operating at a high flow rate. The deformable nature of the nano-sieve enables RBCs retrieval by a simple hydrodynamic deformation. Furthermore, we study the stability of microbeads-stacked micropattern within the nano-sieve channel via fluorescence microscope, optical tomography, microfluidics, and computational fluidic calculation thus building up a reliable system for the successful plasma extraction. By varying the mechanical properties of the nano-sieve, a new model is established to better explain the fluid-structure interaction. This plasma extraction system capable of RBCs retrieval can be widely used for many clinical applications either in hospitals or point-of-care (POC) settings.

### Experimental Details

Materials: Polydimethylsiloxane (PDMS, Sylgard®184) was obtained from Krayden Inc., CO, USA. Glass wafer (D263, thickness: 550 μm) was purchased from University Wafer, MA, USA. Silica magnetic beads (diameter: 10 μm) were received from Alpha Nanotech Inc., Vancouver, Canada. PBS buffer solution (1× without calcium and magnesium, pH 7.4 ± 0.1) was ordered from Corning Inc., NY, USA. Fresh human blood with EDTA was obtained from StemCell Technology Inc., Vancouver, Canada.

PDMS preparation: PDMS curing agent and base were mixed in a weight to weight ratio of 1:5, 1:10, and 1:20, respectively. The PDMS mixture was degassed in a vacuum chamber for 2 hrs before molding. The steel mold was fabricated following the Type IV specimen standard in ASTM D638 (length: 33 mm; width: 6 mm; depth: 6 mm). After pouring the PDMS mixture into the metal mold, it was left in room temperature for 5 hrs to eliminate any air bubbles caused by stirring mixing. The mixture was then cured in an oven at 65°C for 12 hrs. After de-molding, the PDMS replica was left in ambient environment for 24 hrs before mechanical testing.

Poisson’s ratio calculation: Monotonic uniaxial tension test was performed on an Instron 5567 universal testing machine following ASTM D638 standard. Each PDMS was stressed to rupture under three testing speeds of 0.1 mm/s, 0.3 mm/s, and 1 mm/s. A 3-D digital image correlation system (LaVision®, Germany) was used to measure the true strain of the sample. The Poisson’s ratio was obtained by calculating the deformation perpendicular to the loading direction.

Hardness testing: The PDMS sample was subjected to Shore A scale hardness measurement in accordance with the ASTM D2240-15 using a FstDgte Shore A durometer. To assure the constant test rate and parallel contact of the durometer to the specimen, the durometer was mounted on a TCD console. A cuboid sample with a thickness of 10 mm was fabricated by using different mixing ratios. At least 10 different points were tested on a single sample with at least 6 mm apart from each point, as recommended by the ASTM standard.

Nano-sieve device fabrication: A 200 nm tetraethyl orthosilicate (TEOS) was deposited on a glass wafer and the nano-sieve was defined by standard photolithography (length: 10 mm; width: 2 mm). Buffered oxide etching (BOE) was used to etch through the TEOS layer. The channel was filled with positive photoresist (AZ-701) with a thickness of ∼1 µm as a sacrificial layer. A PDMS slab (thickness: 5 mm) was bonded on the TEOS layer by oxygen plasma (Electro-Technic Product, BD-20AC Corona Treater). The device was then baked at 100 °C on a hotplate for 2 hrs. The inlet and outlet were created by a biopsy punch (diameter: 1 mm). Finally, acetone was injected into the nano-sieve channel to completely remove the positive photoresist, followed by DI water cleaning.

Microbeads stacking and characterization: For each experiment, 30 µL of the silica magnetic microbeads (∼ 7.6 × 10^7^ beads/mL) were introduced into the nano-sieve channel by a syringe pump at a flow rate ranging from 20 to 60 µL/min. Fluorescence microscope (AmScope, 10 ×) equipped with a high-speed camera (AxioCam MRc, Zeiss) was used to capture the microbeads pattern in the nano-sieve channel and the outlet reservoir.

Optical coherence tomography and processing: Nano-sieve channels were imaged via an optical coherence tomography (OCT) system, which is a label-free, volumetric imaging modality with an axial resolution of 3 μm and a lateral resolution of 4 μm. The field of view of OCT system is 3 × 6 × 1.94 mm^3^. The imaging system utilizes a raster scanning pattern and works at a line rate of 25 kHz. To evaluate the topology of the nano-sieve channels, volumetric dataset was processed by a K-mean clustering algorithm to segment nano-sieve region from background [28]. From segmented regions, the maximum height and volume were measured and grouped regarding flow rates.

Plasma separation and RBCs retrieval: Before on-chip experiments, whole blood sample containing RBCs was imaged with 20× magnification by a bright-field microscope (AmScope). For plasma separation, whole blood sample was first diluted to 45 µL with PBS buffer and then introduced into the nano-sieve with a flow rate of 5 µL/min. After plasma collection, a high flow rate (∼900 µL/min) was applied to deform the nano-sieve and retrieve the microbeads and RBCs in a new Eppendorf tube. Finally, a neodymium magnet was used to isolate the microbeads and RBCs.

## Results

Our approach of separating human plasma from whole blood sample is presented in **Figure 1a**. The magnetic beads with a diameter of 10 μm were pumped into the nano-sieve channel to construct a 3-D microbeads array. Diluted whole blood was then injected into the microbeads-stacked nano-sieve for the separation process. The deformed PDMS slab holds the microbeads in the nano-sieve and cannot be removed with a flow rate under 60 µL/min. As shown in **Figure 1b**, patterned by standard microfabrication, several nano-sieve channels can be operated simultaneously for multiplexing plasma separation.

**Figure 1.**
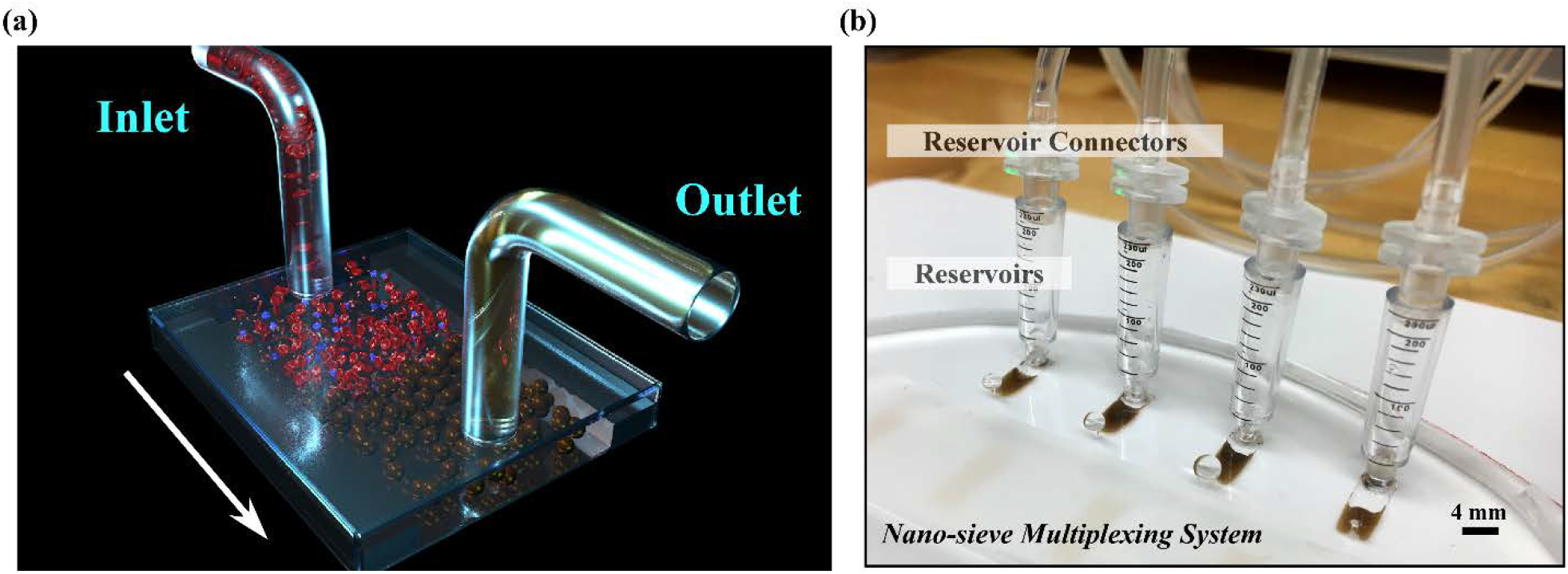
(a) Schematic of the microbeads packed nano-sieve device for the separation of blood cells and plasma. (b) Micrograph of the multiplexing nano-sieve system for plasma separation and RBCs retrieval.

To understand the microbeads patterning process and maximize the patterning capacity, we studied the microbeads patterning with both fluorescence microscope and optical tomography. As shown in **Figure 2a**, for 1:5 PDMS samples, most of the microbeads are contained in the nano-sieve without leaking issues. At a low flow rate, the microbeads are pushed to the sides of the nano-sieve (**Figure 2a-i**). Increasing the flow rate causes the microbeads to accumulate to the middle of the channel (**Figure 2a-ii** and **2a-iii**). Similarly, for 1:10 PDMS samples, the microbeads are distributed to the sides of the nano-sieve at a low flow rate (**Figure 2a-iv**) and tend to accumulate to the middle of the channel at higher flow rates without leaking issues (**Figure 2a-v** and **2a-vi**). However, for 1:20 PDMS samples, irregular patterns and significant microbeads leaking are observed even at a low flow rate (**Figure 2a-vii**). Increasing the flow rate results in the severe microbeads leaking to the outlet reservoir (**Figure 2a-viii** and **2a-ix**). This observation agrees with the volumetric measurements based on OTC scanning. As shown in **Figure 2b** and **2c**, a significant increase of volume and maximum height from 20 µL/min to 40 µL/min are observed for both 1:5 and 1:10 PDMS samples. Further increasing the flow rate to 60 µL/min doesn’t increase the volume and maximum height, agreeing with the microscope images. The measured volume for 1:10 PDMS sample is ∼2 times larger than 1:5 PDMS sample, indicating a larger cross-section plane for fluidic passage. On the other hand, for 1:20 PDMS samples, a distinct drop in microbeads volume at a high flow rate is observed, indicating a significant microbeads loss (**Figure 2d**).

**Figure 2.**
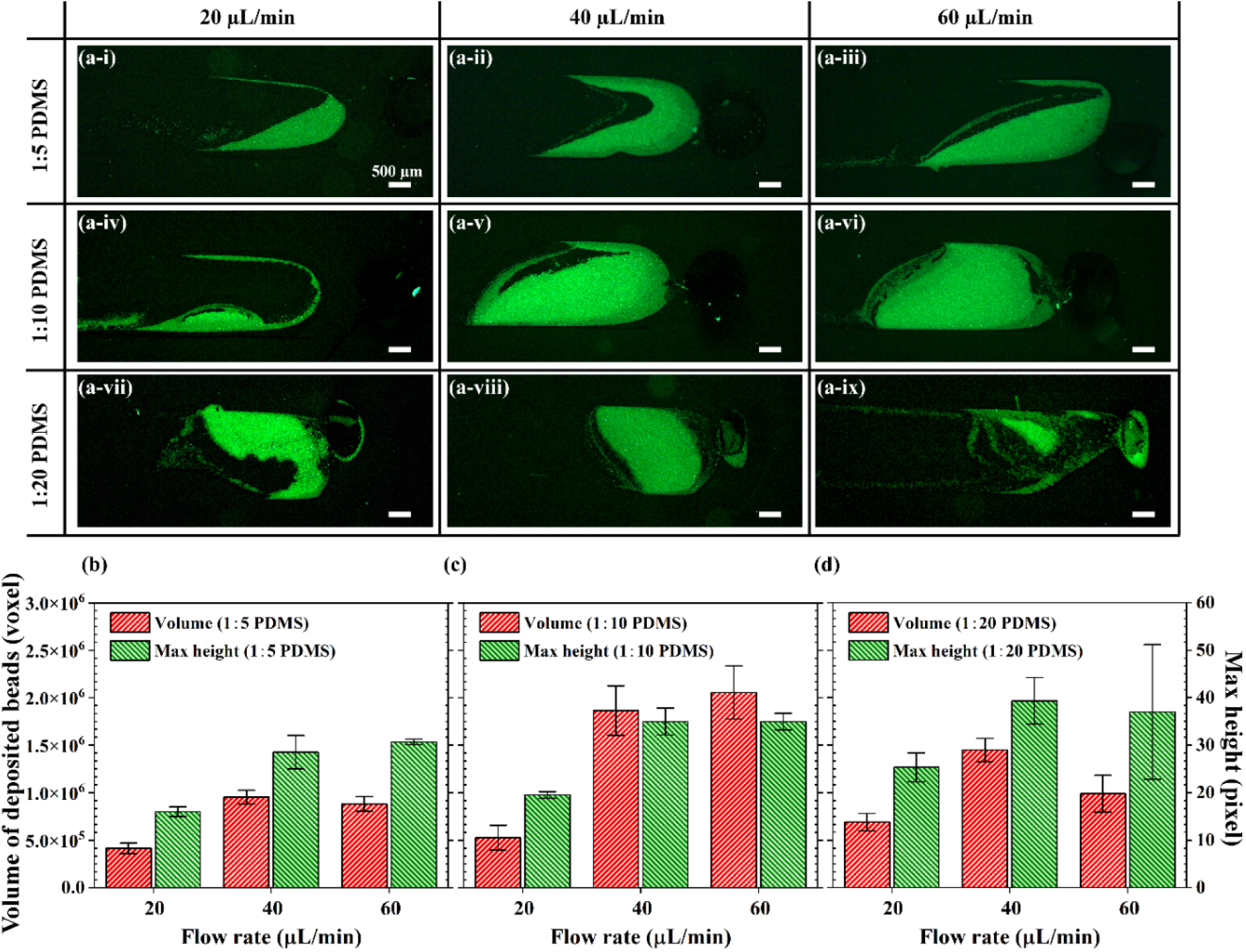
(a) Fluorescence microscope images of the packed microbeads in nano-sieve at various flow rates (20-60 µL/min) and PDMS curing agent/base ratio (1:5 to 1:20). Measured volume and maximum height of the microbeads packed nano-sieve by optical tomography with curing agent/base ratio of: (b)1:5; (c) 1:10; and (d) 1:20.

The measured Passion’s ratio and Shore hardness are shown in **Figure 3a and 3b**, respectively. The Poisson’s ratio versus true strain shows a similar value for all the samples, regardless of the mixing ratio. On the other hand, the measured hardness of 1:10 sample is slightly higher than the 1:5 sample and is ∼2.5 times higher than the 1:20 sample, suggesting a significant drop of mechanical properties when increasing the mixing ratio from 1:10 to 1:20.

**Figure 3.**
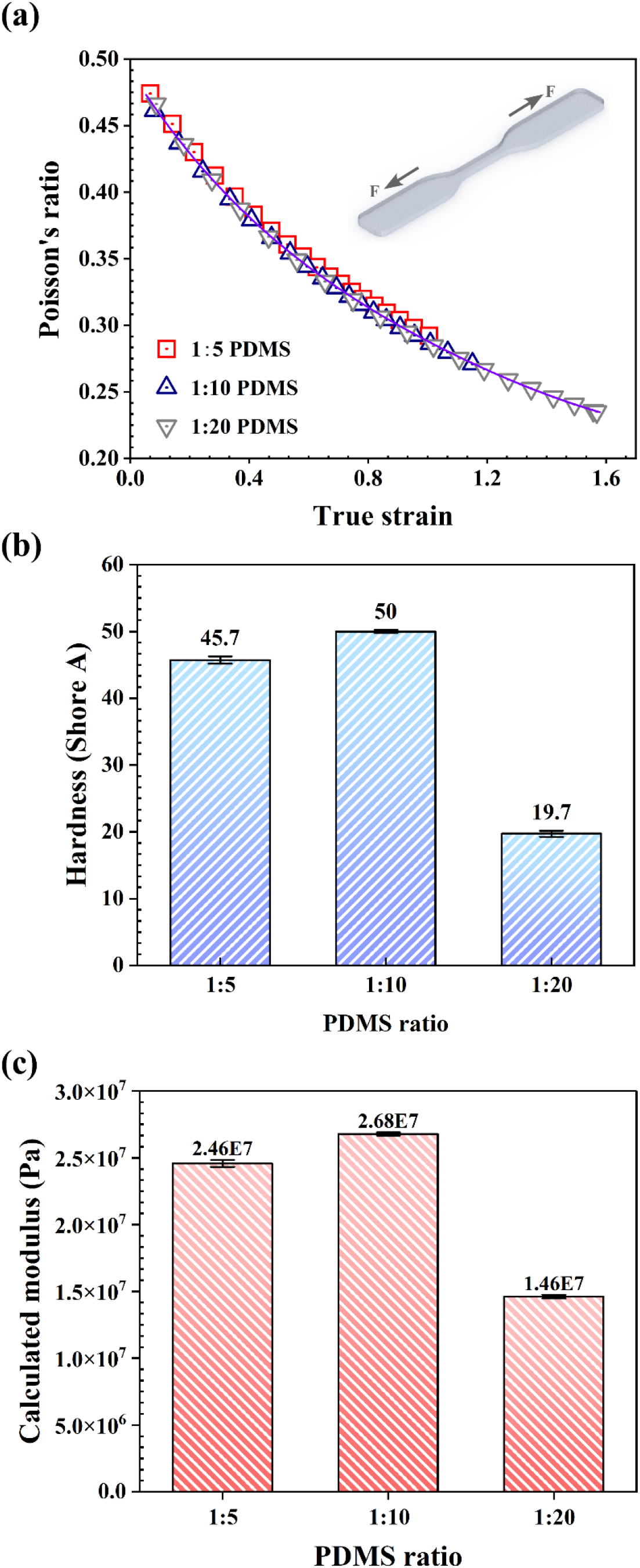
Measured mechanical properties of PDMS with a curing agent/base ratio ranging from 1:5 to 1:20: (a) True stress versus true strain relation. b) Poisson’s ratio with a true strain ranging from 0 to 1.6. The inset shows the specific PDMS model for the mechanical testing. (c) Elastic modulus. (d) Hardness.

Since the deformable nano-sieve device works within a small deformation range, we use the Shore hardness test to estimate the Young’s modulus,

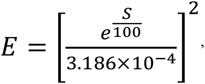

where *S* is the Shore durometer and *E* is the Young’s modulus [29].

With the mechanical properties, we then constructed a theoretical model to understand the microbeads trapping and leaking. The force balance of a single microbead under high flow rate through the channel is shown in **Figure 4a**. The problem is simplified as a microsphere being trapped between two deformable surfaces. Reaction forces normal to the surfaces and toward the particles are applied at the level of

**Figure 4.**
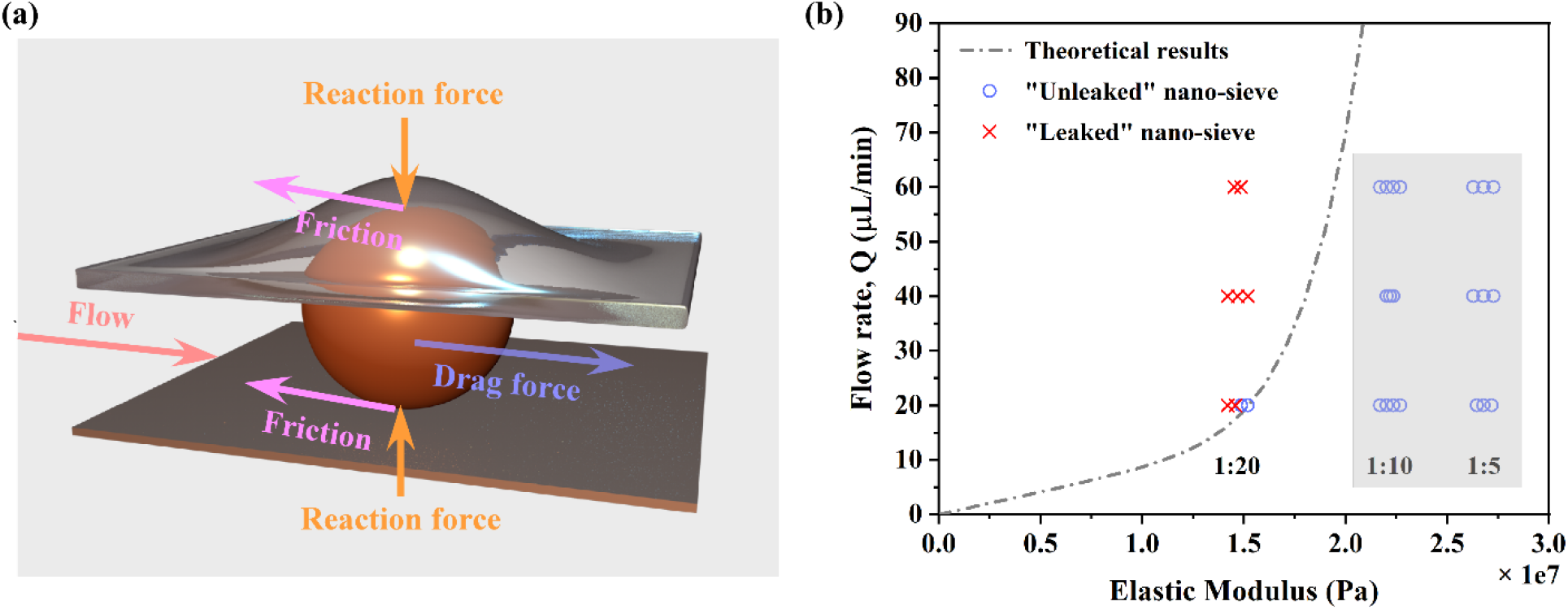
(a) Force analysis of a single microbead sandwiched between two deformable surfaces. (b) “Leaked” (red) versus “unleaked” nano-sieve (blue) at various flow rates observed under a fluorescence microscope. The grey dashed line is the “leaking” boundary based on the theoretical prediction, agreeing with the experimental results.

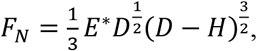

where *F*_*N*_ is the reaction force in the normal direction

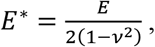

*E* is the elastic modulus, *v* is the Poisson’s ratio, *D* is the microbead diameter, and *H* is the original height of the gap between two surfaces. The maximum possible friction force, *F*_*R*_, is obtained using the formula

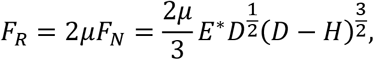

where *μ* is the coefficient of friction. While the flow drags the sphere toward to the outlet, the drag force, *F*_*D*_, is modeled using

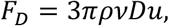

where *ρ* is the density of the fluid, *υ* is the fluid viscosity, and the cross-section average flow velocity is

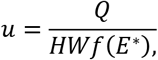

which is a function of the total discharge *Q*, the gap height *H* before deformation, the gap width *W*, and the elastic modulus *E*. The function *f(E*^*\**^), indicating the ratio of the deformed cross-sectional area to the original, is incorporated to take account of the deformation of the channel. As the two extremes, if the Young’s modulus is infinitely large, the gap will be enlarged to the uniform height of particle diameter *D*, while if the material is infinitely soft, the gap will be the original height of *H*. Here, we use a fitting model,

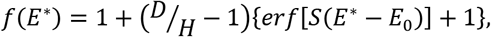

to capture the trend that the average gap height increases with greater elastic modulus transitioning from *H* to *D*. The error function is used to capture the transition with a fitting coefficient *S* and *E*_*0*_ to match the deformation rate and flexibility. In this theoretical model, we thus hypothesize that microbeads will be leaked if the drag force is greater than the maximum friction (*F*_*D*_ *> F*_*R*_) and no leakage occurs if the drag is less than the highest possible friction (*F*_*D*_ *> F*_*R*_). With the fitting coefficients of *S* = 10^−7^ and *E*_*0*_ = 3.3 × 10^7^ Pa, the theoretical model can precisely predict the boundary (dashed line) between leakage and no leakage cases, as shown in **Figure 4b** (“×”: leakage; “**○**”: no leakage). In addition, the current model well explains the mixed results observed for 1:20 PDMS sample at a flow rate of 20 µL/min.

As the 1:10 PDMS sample provides the largest volume for microbeads packing without leaking issues, we then use this sample for human plasma separation and RBCs retrieval. A micrograph of the separation process is shown in **Figure 5a**. The whole blood with a red color enters the nano-sieve and the RBCs are trapped by the microbeads array, allowing the human plasma passing through the voids of the microbeads. The separated human plasma has a light-yellow color, as shown in the outlet tubing. **Figure 5b** shows the RBCs clogged nano-sieve under a bright-field microscope. The blood cells are randomly mixed with the microbeads in the nano-sieve and most of them are blocked at the microfluidic inlet. To characterize the plasma separation efficiency, the original blood sample and the filtered sample are shown in **Figure 5c-i** and **5c-ii**, respectively. Original sample has a high concentration of RBCs. On the other hand, the filtered sample shows only a few individual cells, indicating a high plasma separation capability. By applying a high flow rate of 900 µL/min, the trapped RBCs were efficiently retrieved in a new Eppendorf tube with a cell density of ∼0.0030 cells/μm^2^, comparable to the original sample (∼0.0026 cells/μm^2^), as shown in **Figure 5c-i** and **5c-iii**.

**Figure 5.**
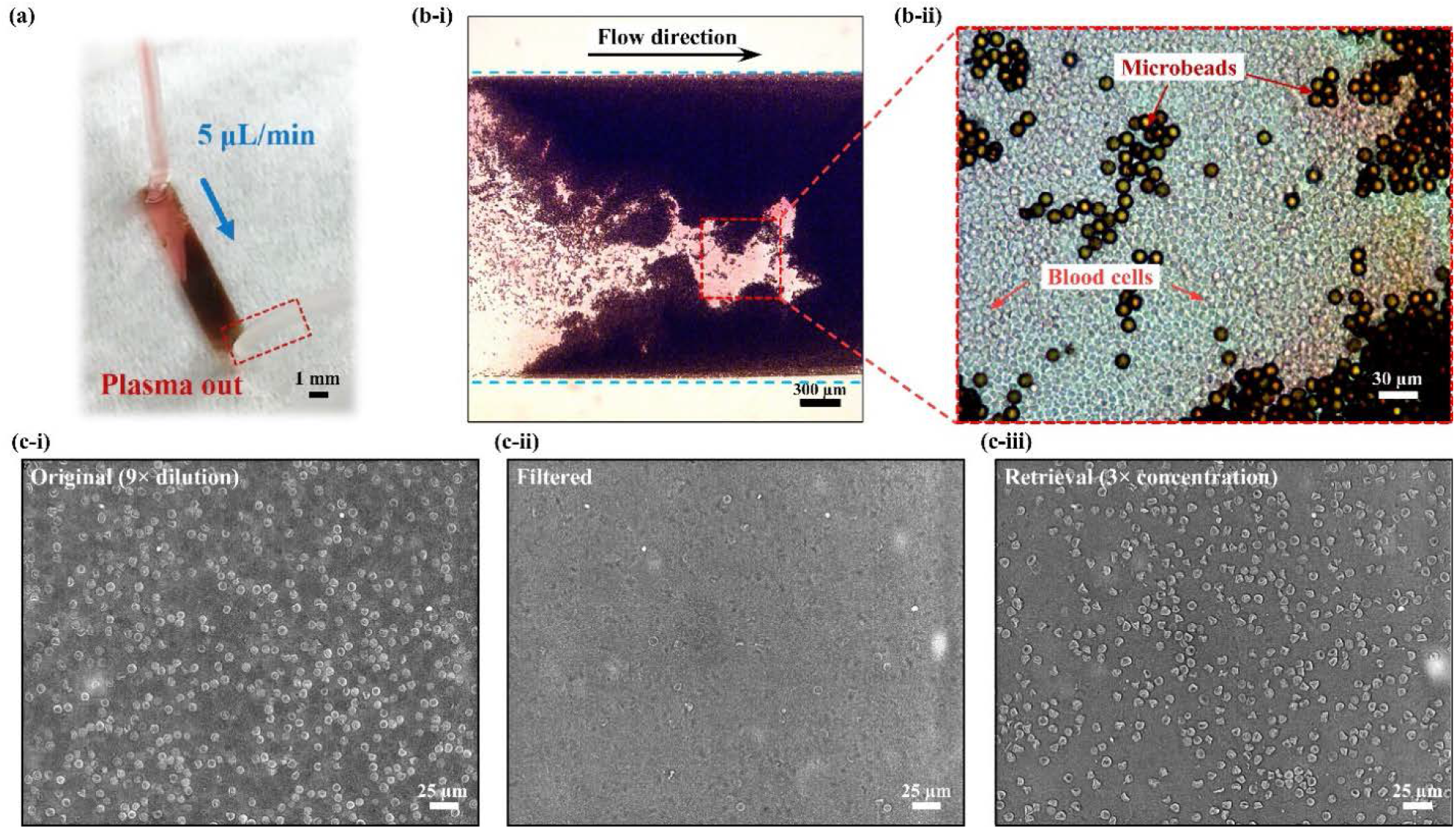
(a) Optical micrograph of a working nano-sieve for the isolation of plasma. The color of the fluids at the outlet is more transparent as blood cells are removed. (b) Bright-field microscope image showing the microbeads pattern and captured RBCs: (i) low magnification; (ii) high magnification. (c) Microscope image of (i) original whole blood showing a high concentration of RBCs; (ii) Filtered plasma solution without RBCs. (iii) Retrieved RBCs from the nano-sieve.

## Discussion

In this work, we experimentally measure the mechanical properties of PDMS with various mixing ratios. Combining with optical tomography scanning, we develop a computational fluidic system to understand how the mechanical properties of the deformable microfluidics and applied flow rate can affect the microbeads packing and leaking. This model successfully explains how the microbeads matrix are formed in the microfluidic channel, enhancing the overall performance of nano-sieve’s plasma separation efficiency.

The efficient plasma separation [30,31] is critical for numerous down-stream molecular analysis and diagnosis [7,32–34]. The microbeads packed nano-sieve device is capable of plasma separation without using bulky and complicated instruments such as a centrifuge. This physical filtering process is easily operated by directly injecting the blood sample into the nano-sieve. Even though immunoassay-based RBCs removal strategy also demonstrates a high plasma separation efficiency [35], reagents such as antibodies are delicate and can degrade rapidly, making the device storage challenging in low resource settings. Our separation process solely relies on the silica microbeads stacking in a deformable microfluidic device and can be stored in ambient environment for a very long period of time, ideal for the sample preparation in POC settings. More importantly, ∼100 individual nano-sieve devices can be patterned on a 150 mm wafer scale for multiplexing plasma separation, which is hard to achieve by membrane-based microfluidic devices [36].

We show that RBCs can be retrieved by simply increasing the flow rate and the deformation of the nano-sieve, which is more superior than microstructure-based separation as the clogged RBCs cannot easily be removed from the functional structure [37–39]. The study of RBCs in patient sample can be used to understand and treat many bloodborne diseases such as sepsis [8,9], sickle cell anemia [10,11], and COVID-19 [40,41]. In the future, the geometry of each nano-sieve can be modulated to process a larger volume of blood sample for low concentration pathogen detection. Changing the size and spacing of the microbeads can also be used to separate and concentrate various types of cells and biomarkers. For example, by varying the size of the packed microbeads, it is possible to separate leukocytes from other two main components (erythrocytes and platelets) in whole blood due to the different size and deformability [42]. Furthermore, as the materials properties of the microbeads can be tuned, the nano-sieve can be used to study various cell mechanics problems [43,44] with real-time microscopy imaging in a confined nano-environment.

Furthermore, our study is a meaningful attempt to test the fluid-structure interaction theory in the application of microfluidics from the viewpoint of the structure material properties. The developed theoretical model can be used to guide the design of deformable microfluidics and the good fitting equation will inspire future more detailed model development.

